# Molecular mimicry in deoxy-nucleotide catalysis: the structure of *Escherichia coli* dGTPase reveals the molecular basis of dGTP selectivity: New structural methods offer insight on dGTPases

**DOI:** 10.1101/385401

**Authors:** Christopher O. Barnes, Ying Wu, Jinhu Song, Guowu Lin, Elizabeth L. Baxter, Aaron S. Brewster, Veeranagu Nagarajan, Andrew Holmes, Michael Soltis, Nicholas K. Sauter, Jinwoo Ahn, Aina E. Cohen, Guillermo Calero

**Author notes:** Co-correspondence should be addressed to GC and AEC ( and). These authors contributed equally to this work (Christopher O. Barnes, Ying Wu).

## Abstract

Deoxynucleotide triphosphate triphosphyohydrolyases (dNTPases) play a critical role in cellular survival and DNA replication through the proper maintenance of cellular dNTP pools by hydrolyzing dNTPs into deoxynucleosides and inorganic triphosphate (PPPi). While the vast majority of these enzymes display broad activity towards canonical dNTPs, exemplified by Sterile Alpha Motif (SAM) and Histidine-aspartate (HD) domain-containing protein 1 (SAMHD1), which blocks reverse transcription of retroviruses in macrophages by maintaining dNTP pools at low levels, *Escherichia coli (Ec)-*dGTPase is the only known enzyme that specifically hydrolyzes dGTP. However, the mechanism behind dGTP selectivity is unclear. Here we present the free-, ligand (dGTP)- and inhibitor (GTP)-bound structures of hexameric E. coli dGTPase. To obtain these structures, we applied UV-fluorescence microscopy, video analysis and highly automated goniometer-based instrumentation to map and rapidly position individual crystals randomly-located on fixed target holders, resulting in the highest indexing-rates observed for a serial femtosecond crystallography (SFX) experiment. The structure features a highly dynamic active site where conformational changes are coupled to substrate (dGTP), but not inhibitor binding, since GTP locks dGTPase in its apo form. Moreover, despite no sequence homology, dGTPase and SAMHD1 share similar active site and HD motif architectures; however, dGTPase residues at the end of the substrate-binding pocket mimic Watson Crick interactions providing Guanine base specificity, while a 7 Å cleft separates SAMHD1 residues from dNTP bases, abolishing nucleotide-type discrimination. Furthermore, the structures sheds light into the mechanism by which long distance binding (25 Å) of single stranded DNA in an allosteric site primes the active site by conformationally “opening” a tyrosine gate allowing enhanced substrate binding.

**Significance Statement:** dNTPases play a critical role in cellular survival through maintenance of cellular dNTP. While dNTPases display activity towards dNTPs, such as SAMHD1 –which blocks reverse transcription of HIV-1 in macrophages– Escherichia coli (Ec)-dGTPase is the only known enzyme that specifically hydrolyzes dGTP. Here we use novel free electron laser data collection to shed light into the mechanisms of (Ec)-dGTPase selectivity. The structure features a dynamic active site where conformational changes are coupled to dGTP binding. Moreover, despite no sequence homology between (Ec)-dGTPase and SAMHD1, both enzymes share similar active site architectures; however, dGTPase residues at the end of the substrate-binding pocket provide dGTP specificity, while a 7 Å cleft separates SAMHD1 residues from dNTP.

## Introduction

Cellular regulation of deoxynucleotide triphosphate (dNTP) pools is a vital process for DNA replication and survival in both prokaryotic and eukaryotic cells. It is tightly controlled by deoxynucleotide triphosphohydrolases (dNTPases) (1-6) through hydrolyzing dNTPs into deoxynucleosides and inorganic triphosphate (PPP_i_) (2, 7-12), As a class of metalloenzymes, dNTPases contain a histidine-aspartate (HD) motif, which helps coordinate a divalent cation near the active site to promote phosphohydrolase activity (13). Given the importance of intracellular dNTP concentrations for cellular survival, studies have shown that tight control of dNTP pools by dNTPases acts as a host mechanism in the cellular defense against pathogens. For instance, in primates the Sterile Alpha Motif (SAM) and Histidine-aspartate (HD) domain-containing protein 1 (SAMHD1) (14) blocks reverse transcription of retroviruses (e.g. HIV-1, SIV) by maintaining dNTP pools at low levels in infected cells (7-10, 12, 15). In addition, observations in prokaryotes have revealed a dNTPase-pathogen interplay, illustrated by *Escherichia coli* (*Ec*) dGTPase, where the gene 1.2 product encoded by bacteriophage T7 inhibits *Ec-*dGTPase to promote productive infection (16).

Crystal structures of the broadly acting tetrameric dNTPases, exemplified by SAMHD1, have enabled a better understanding of phosphohydrolase activity by revealing the binding modes of activator or allosteric regulator nucleotides, as well as dNTP substrates (10-12, 17). However, unlike the vast majority of dNTPases displaying broad activity towards canonical and non-canonical dNTPs (11, 12, 15, 18, 19), *Ec-*dGTPase displays a strong preference for hydrolyzing only dGTP substrates (2, 10, 11, 15, 17, 19, 20). The structural basis of *Ec-*dGTPase specificity towards dGTP is lesser understood despite recent crystal structures of a free and ssDNA bound *Ec*-dGTPase (21). Indeed, the potential of ssDNA cofactors effecting the dGTP binding site (21, 22), suggests a unique mechanism significantly different in the allosteric activation of phosphohydrolase activity when compared to other dNTPase enzymes.

In the present work, we aimed to understand the structural basis of *Ec-*dGTPase substrate specificity and the mechanism of phosphohydrolase activity in the presence of nucleotide substrates. Given the importance of the metal cofactor in *Ec*-dGTPase activity (2, 13, 19), we determined the radiation-damage free crystal structure in the presence of Mn^2+^ cations to 3.2 Å using data collected at the Linac Coherent Light Source (LCLS) X-ray Free Electron Laser (XFEL). These experiments employed a new methodology for highly efficient serial diffraction (SFX) experiments using micrometer-sized crystals that builds upon conventional crystal visualization and fixed target diffraction techniques (23, 24). Our methodology combines 1) the use of specialized multi-crystal holders (MCH) compatible with cryo-protectant or native crystallization conditions, 2) the identification of individual micro-crystals on MCHs through the analysis UV-microscopy images, and 3) the development of automated routines for serial positioning of the identified crystals during data collection. Implementation of these methods improved on similar reference and alignment schemes (25-31) enabling rapid mapping and positioning of multiple crystals in random locations on the holder, which consistently resulted in >80% crystal hit-rates and the highest indexing rates reported to date for any SFX experiment.

In addition, we investigated substrate specificity by applying chemical cross-linking methods to introduce nucleotide substrates into the catalytic site of *Ec-*dGTPase crystals, and obtained dGTP-, dGTP-1-thiol-, and GTP-bound *Ec-*dGTPase structures. We found that structural elements from an adjacent monomer, which protrude into the enzymatic active site, are responsible for nucleotide discrimination and dGTP specificity. Analysis of catalytic residues and Mn^2+^ cations of free- and bound- *Ec-*dGTPase structures, revealed a regulatory mechanism for Tyr^272^ that may explain activation or inhibition by ssDNA and GTP, respectively. Overall, these structures provide detailed mechanistic insights into *Ec-*dGTPase function and demonstrate a conserved binding mode for nucleotide substrates across dNTPase enzymes.

## Results

### XFEL data collection of microcrystals on Multi-Crystal Holders (MCHs)

To address the scarcity of XFEL beam time, we sought to improve the efficiency and reliability of data collection methods by simplifying related sample preparation and operational requirements for fixed-target setups. To this end, we developed a Multi-Crystal Holder (MCH) (Fig. S1A, B) capable of holding microcrystals under native- or cryo- crystallization conditions that is compatible with UV fluorescence microscopy for crystal imaging and identification (Fig. 1 and Fig. S1, see Methods). Given the high contrast of the UV fluorescence signal of crystals versus the solvent background, bright areas corresponding to the location of individual crystals on MCHs were identified and mapped relative to four fiducial marks (Fig. 1B and Fig. S1). Algorithms previously incorporated into the Blu-Ice/DCSS experimental control system to position grid-based sample holders and microfluidic traps at LCLS-XPP (23, 26, 27) were adapted to automatically position crystals based on their relationship to the four fiducial coordinates of the MCH for efficient serial diffraction experiments (Fig. 1B, Fig. S1D, E, and Table S1). This protocol was applied to various crystal morphologies generated from *Ec-*dGTPase or Pol II complexes mounted on MCHs (Fig. S1), enabling diffraction data to be collected in an automated fashion with improved intensities for high angle Bragg reflections, consistent with previous results using MCHs (32, 33).

**Figure 1.**
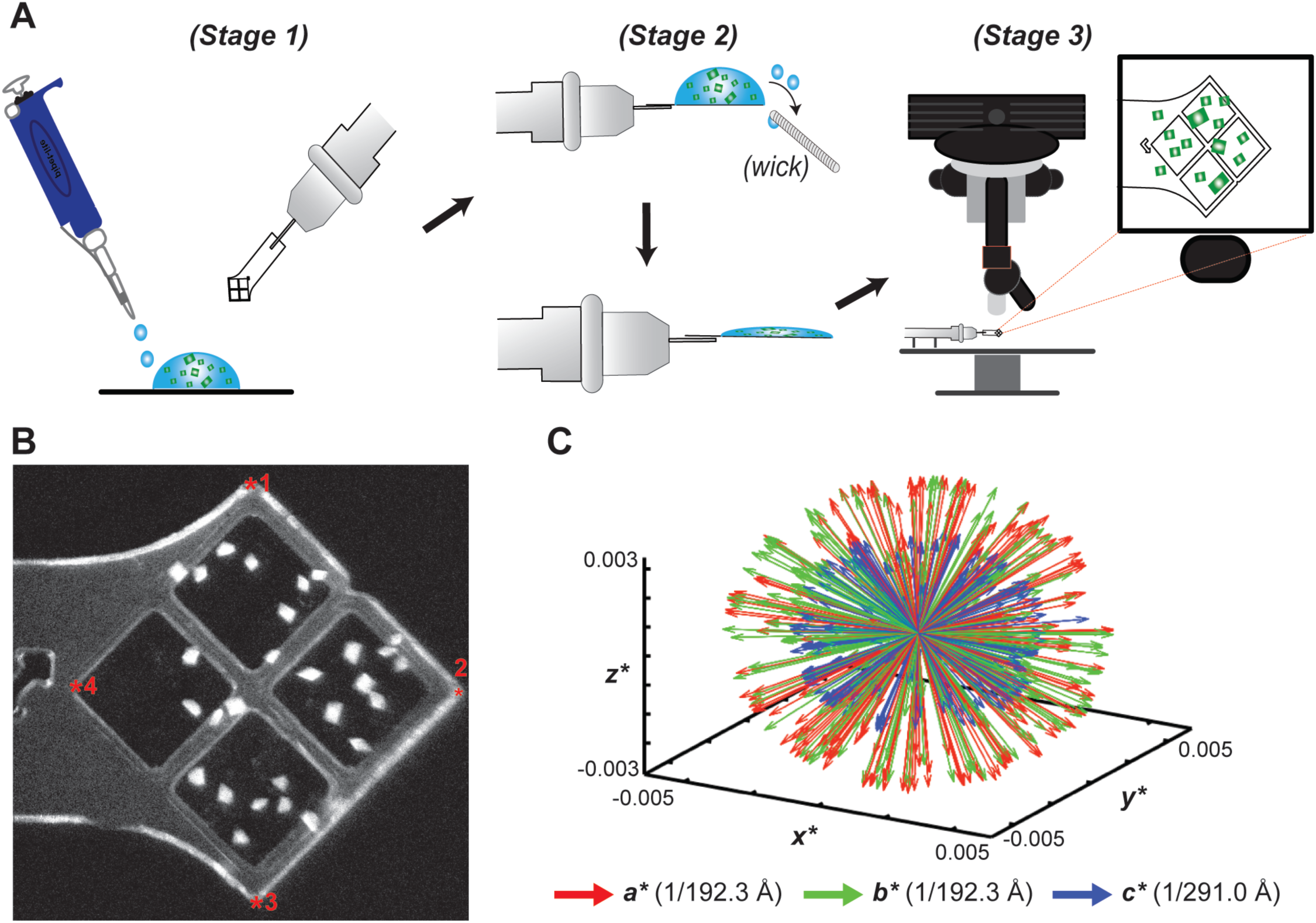
Loading and mapping of crystals on MCHs. **A)** Schematic representation of MCH loading as detailed in the experimental methods section**, B)** UV microscopy image of Pol-Spt4/5 mounted crystals. Fiducial marks are illustrated as red asterisk**. C)** Reciprocal space representation of the basis vectors of 221 indexed dGTPase images demonstrating the lack of preferential alignment when mounting in MCHs.

In addition, we implemented multi-shot data collection strategies for crystals measuring >100 microns along a single axis (Fig. S1I-K). Similar to helical data collection protocols at LCLS (23, 24, 34), translation of approximately 50 µm into uncompromised crystal volumes with angular offsets from the origin were introduced to increase dataset completeness. To test if micro-crystals displayed problems of preferential orientation on the MCH surface, we analyzed the reciprocal basis vectors for indexed *Ec-*dGTPase and Pol II crystals that were singly exposed for one MCH (Fig. S2) and for the entire *Ec-*dGTPase dataset (Fig. 1D), using described methods (35). Our results revealed a spherical-like projection indicative of randomly oriented crystals on MCHs (23, 24, 26, 34). As a result, all datasets collected showed >90% completeness to high-resolution, with only 221 still images being required to solve the structure of the *Ec-*dGTPase enzyme (Table S2). Overall, application of MCH methodologies resulted in efficient data collection with minimal background scattering to preserve weak Bragg reflections, highly accurate microcrystal hit rates (>80%), increased indexing rates, and dataset completeness from a minimal number of exposed crystals (Table S2).

### Ec-dGTPase structure and active site metal coordination

The radiation damage-free XFEL structure of the *E*c-dGTPase apo-enzyme to 3.2 Å was solved from <150 crystals exposed to the XFEL beam (Fig. 2), representing the fewest number of randomly oriented micro-crystals used to obtain a complete SFX dataset. Initial phases were generated by molecular replacement using a selenium-methionine SAD phased structure (Fig. S2A) as a search model (Table 1). The hexameric *Ec-*dGTPase XFEL structure (Fig. 2A) is consistent with previously published dGTPase apo-structures from *Pseudomona syringae* (PDB:ID 2PGS), *E. coli* (PDB:ID 4XDS), and the *E. coli* apo-structure bound to single stranded DNA in which a slight conformational change occurred in the *α*10 helix of the substrate binding pocket (ssDNA, PDB:ID 4×9E) (Fig. S2C).

**Table 1.**
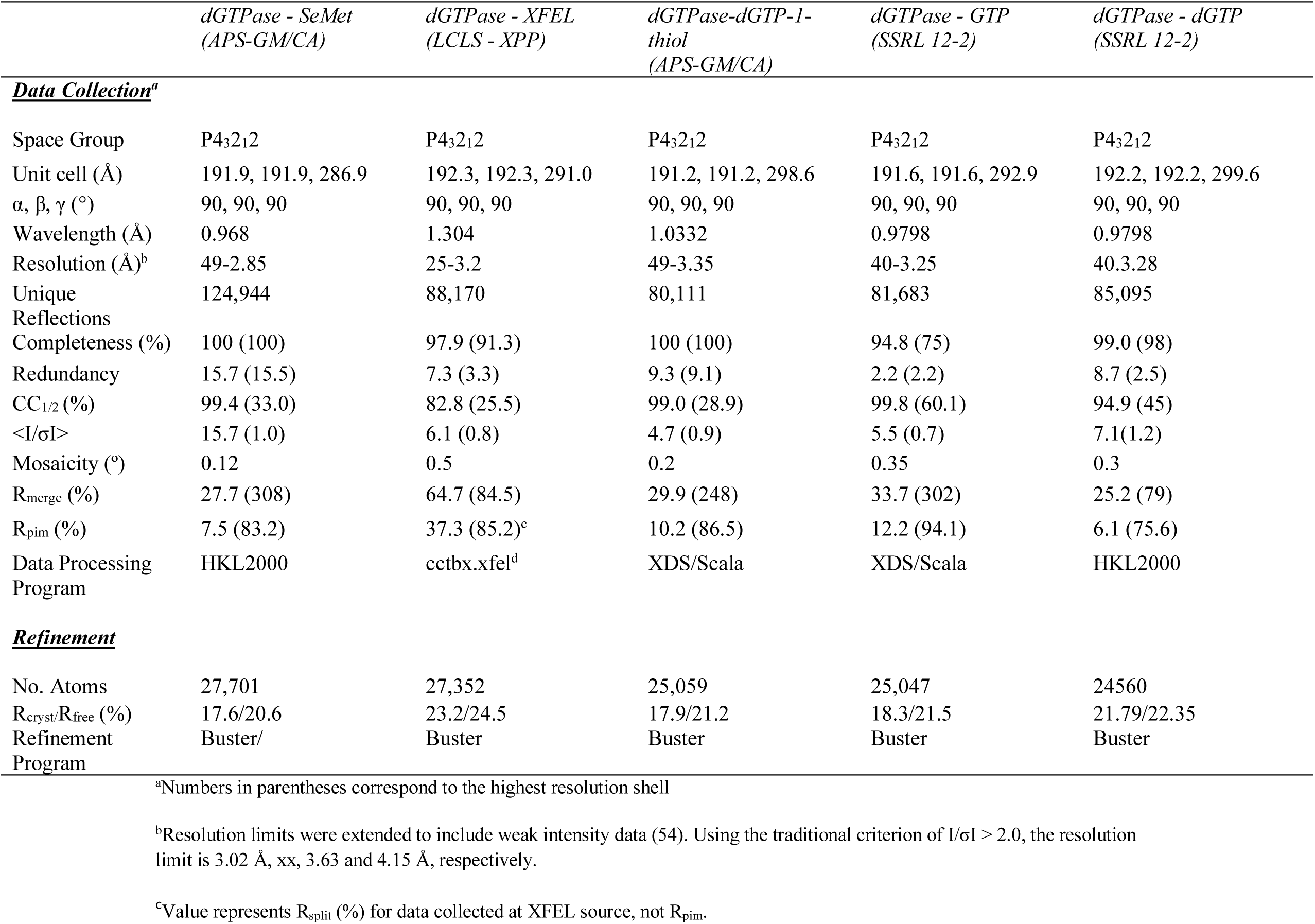
Crystallographic data and refinement statistics

**Figure 2.**
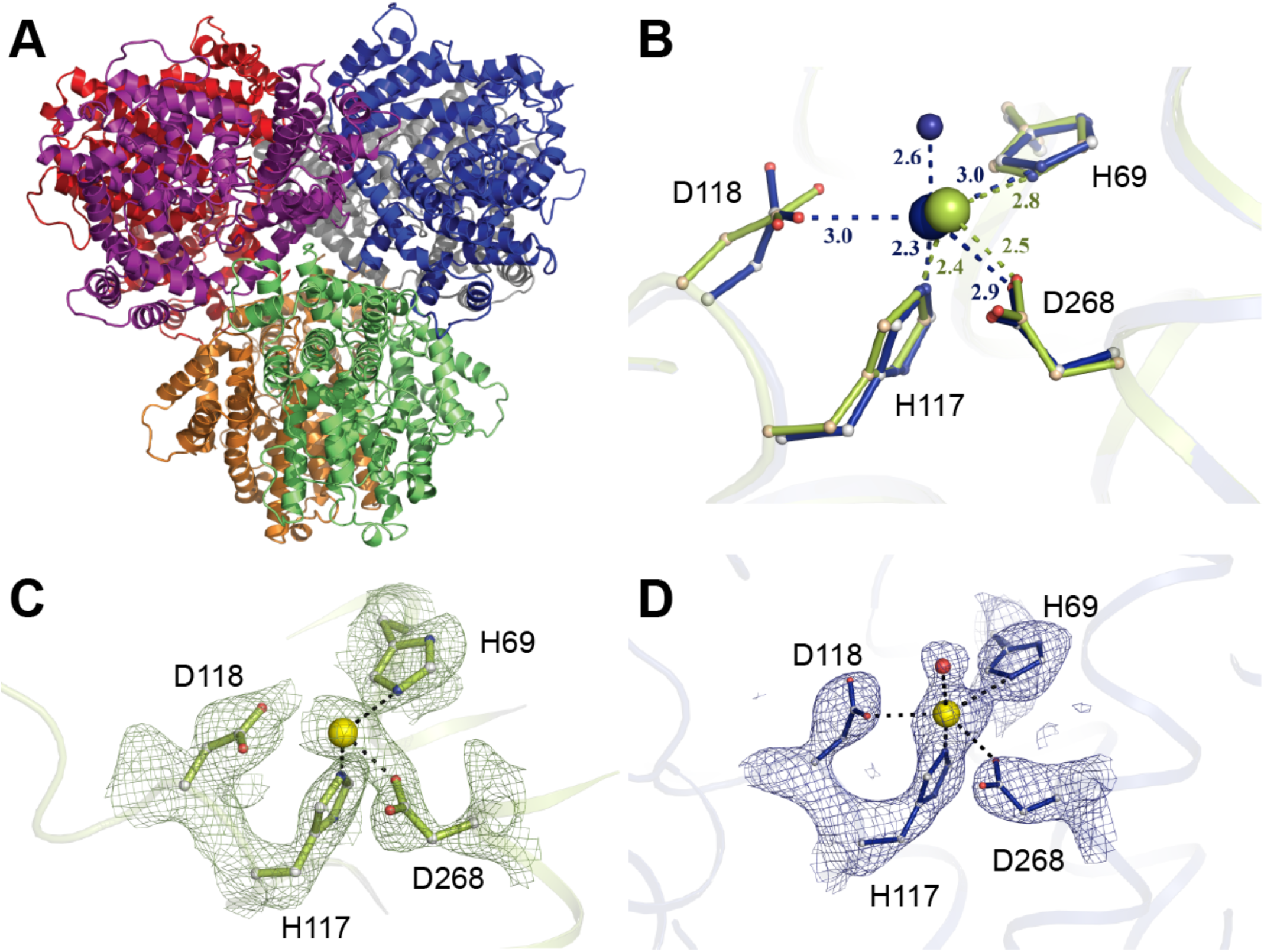
The hexameric *Ec-*dGTPase XFEL crystal structure. **A)** Cartoon representation of the hexameric *Ec-*dGTPase XFEL crystal structure solved using 221 still images from randomly oriented crystals**. B)** Differences in Mn^2+^ coordination for XFEL (blue) and low-dose synchrotron structure (shown in olive green). Hereafter, potential H-bond interactions with distances shorter than 3.5 Å are indicated as dashes between residues. **C)** Electron density of the Sigma-A weighted *2F*_*obs*_ *– F*_*calc*_ map contoured at 1.5*σ* for residues comprising the HD motif (synchrotron data); Mn^2+^ ion is illustrated as a yellow sphere. **D)** Electron density of the Sigma-A weighted *2F*_*obs*_ *– F*_*calc*_ map contoured at 1.5*σ* for residues comprising the HD motif (XFEL data). A water molecule (indicated in red) is seen to form part of Mn^2+^ coordination.

However, comparison of conserved residues in the HD-motif (His^69^, His^117^, Asp^118^, and Asp^268^) revealed subtle changes in metal coordination at the catalytic site. The initial unbiased sigma A-weighted difference map (F_0_-F_calc_) of the radiation damage-free XFEL structure showed electron density for the Mn^2+^ ion, the HD motif residues and a previously unobserved water molecule that contributes to Mn^2+^ coordination (Fig. 2B and Fig. S2B). Moreover, Asp^118^ was on average 0.5 – 0.7 Å closer to the Mn^2+^ ion across all six monomers when compared to the Se-Met apo-structure, which had an averaged absorbed X-ray dose of 3.4 MGy (Fig. 2C, D). This positional difference places the catalytic metal within its coordination sphere and suggests that at even fairly low X-ray doses, radiation damage is accrued site-specifically around metal centers, possibly through the decarboxylation of the neighboring aspartic acid residue (36).

### Structural basis for dGTP binding and phosphohydrolase activity

Next, we wanted to characterize the substrate bound form of *Ec-*dGTPase by determining the structure in the presence of dGTP and the non-hydrolysable dGTP analog, dGTP-1-thiol. Initial co-crystallization experiments of *Ec-*dGTPase with Mn^2+^ ions and dGTP-1-thiol revealed no ligand-bound structures, likely due to the high-salt condition (1.8 M ammonium sulfate) from which the crystals were harvested. Thus, we employed chemical cross-linking using glutaraldehyde to stabilize the integrity of the crystal-lattice, while decreasing the salt concentrations to physiological levels for overnight soaking experiments with dGTP or dGTP-1 thiol (see methods). Crystals remained stable during this procedure resulting in conventional synchrotron-based structures with bound substrates and metals in the active site (Fig. S3A-E). Cross-linked crystals also showed catalytic activity when incubated with dGTP, suggesting that glutaraldehyde had no adverse effects on phosphohydrolase activity (Fig. S3B).

Comparisons between the substrate-bound and apo-structures show a root mean square deviation (RMSD) of 0.9 Å for the catalytic versus 0.3 Å for the non-catalytic regions of the protein. These differences are a result of conformational changes induced by dGTP binding, as active site residues rearrange to provide a tight-fitting pocket for the substrate (Fig. 3A, B). Critical residues involved in dGTP binding, as well as dGTP-1-thiol binding, include: i) π-π stacking interactions between the electron-rich aromatic ring of Phe^391^ and the positively charged guanosine ring; ii) hydrogen bonding between Arg^433^, Glu^400^ and Val^54^ and the guanosine ring; iii) stacking interactions between Tyr^272^ (a highly conserved tyrosine found within dNTP triphosphohydrolases (15, 19, 37) with the sugar pucker; iv) 3′-OH discrimination via Gln^53^ and Asp^276^; v) interactions of the *α*-phosphate with Mn^2+^; and vi) interactions with the *β*- and γ-phosphates through Asn^186^, Lys^211^, Tyr^212^ and Lys^232^ (Fig. 3C,D and Fig. S3C-E). The full icosahedral coordination of the Mn^2+^ ion was also visualized in the substrate-bound structure, comprising the four residues of the HD motif, a water molecule (W_1_) and an oxygen from the dGTP *α*-phosphate that replaces the coordinated water observed in the apo-XFEL structure (Fig 3D). Moreover, among the six hexamers, His^126^ was found as two conformers; the first one occupying a similar position as the substrate-free structure, and the second one positioned in-line with the hydroxyl group of the dGTP α-phosphate (Fig. 3A). This latter conformation is consistent with the proposed nucleophilic substitution reaction mechanism for dNTP hydrolysis via a nearby water molecule (35).

**Figure 3.**
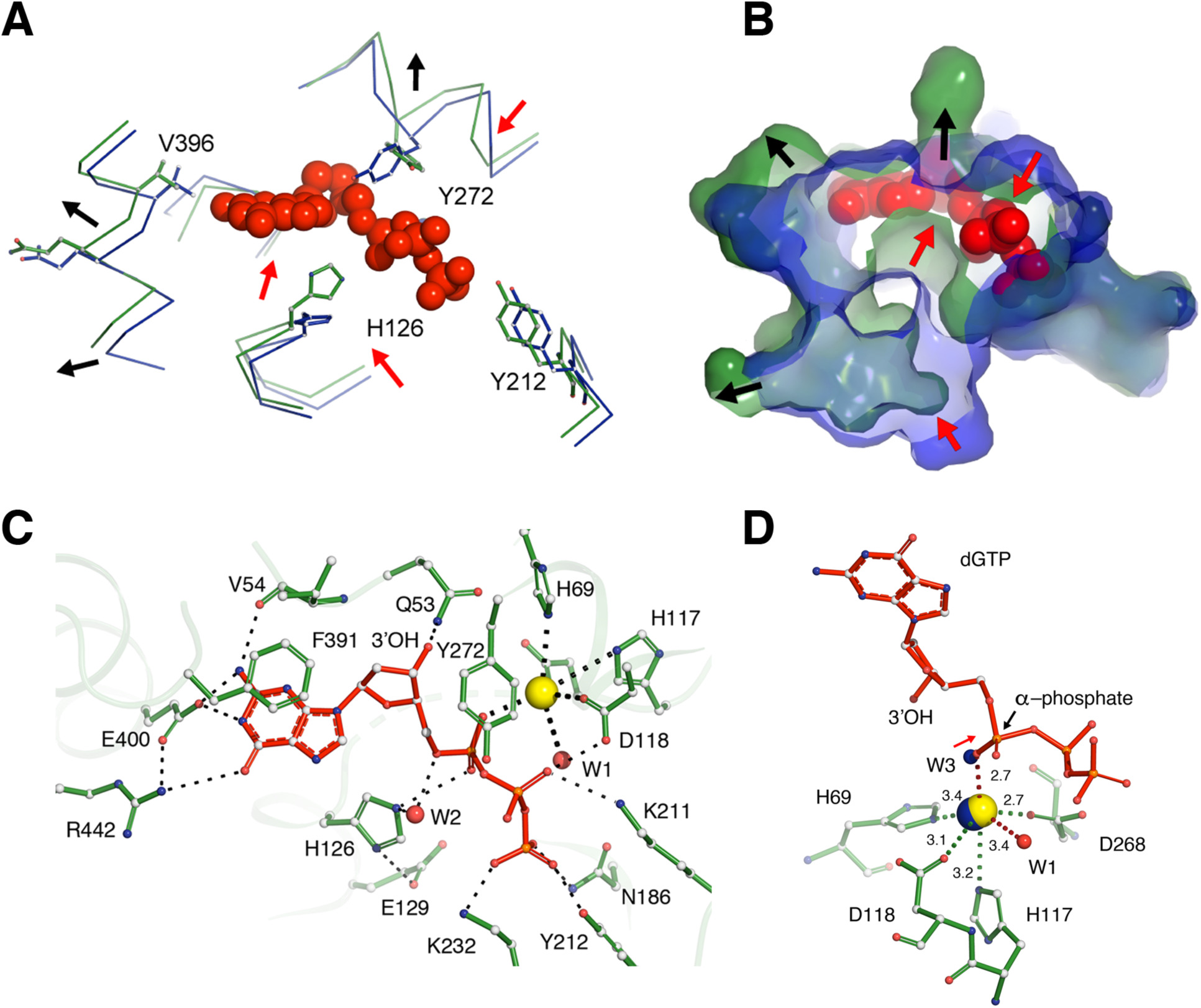
Interactions of dGTP substrate with *Ec-*dGTPase. Wire **(A)** and surface **(B)** representations of the overlay between the apo-(blue) and dGTP-bound (forest green) structures showing the conformational changes observed in the active site pocket upon dGTP binding (red spheres). Red arrows indicate contraction and black arrows indicate expansion of the pocket. **C)** Ball and stick representation of key residues (forest green) involved in dGTP (red) binding. Hydrogen bonds are illustrated as dashes, water molecules as red spheres. **D)** Ball and stick representation of Mn^2+^ (yellow sphere) coordination by dGTP residues (forest green). Distances are indicated next to dashes. The position of the apo-XFEL Mn^2+^ ion and coordinating water after overlay with the dGTP bound-structure are indicated as blue spheres (see text).

Owing to the contrast in side chain confirmation for free- and substrate-bound structures, as well as the observed “sandwiching” between Tyr^272^, His^126^ and the dGTP ligand (Fig. 3C), we effectively mutated these residues to alanine to identify their roles in phosphohydrolase activity. The single alanine mutation in the active site had no effect on expression, purification, or hexamer formation for the H126A, E129A, or Y272A variants when compared to wild-type. Thus, we assayed enzymatic activity by monitoring deoxyguanosine product formation using reverse-phase chromatography. All three variants had abolished hydrolytic activity when compared to wild-type enzyme, even after 2-hour incubation in the presence of 100 µM dGTP and 5 mM MgCl_2_ (Fig. S3G). While the roles of His^126^ and Glu^129^ in catalysis have been well-established (10, 37), the function of Tyr272 is less clear. Similar to structures of the tetrameric dNTPases (PDB 3IRH, PDB 4TNQ, PDB 2DQB (9, 11, 37), His^126^ and Glu^129^ form a catalytic dyad, by which Glu^129^ plays a role in positioning His^126^ in-line with the α-phosphate during catalysis. Interestingly, while mutations in the catalytic dyad showed slight product formation at the two-hour time point, the Y272A variant resulted in a consistently inactive enzyme (Fig. S3G). This suggests that Tyr^272^ plays a critical role in *Ec-*dGTPase phoshydrolase activity, possibly by stabilizing the *α*-phosphate for nucleophilic attack.

### Chracterization of Ec-dGTPase specificity and nucleotide discrimination

Since *Ec-*dGTPase shows a very high affinity and preference for dGTP substrates (2, 19), we determined the mode by which the enzyme can specifically recognize this substrate compared to other purine or pyrimidine rings, namely dTTP, dATP and dCTP. Recognition of dGTP within the active site is achieved by Glu^400^ and the backbone carbonyl of Val^54^, which interacts with the purine amine groups. In addition, residues Arg^433^ and Arg^442^ project into the binding pocket from an adjacent monomer to interact with the ketone group of the purine ring (Fig. 3C and Fig. S3E). These residues create a tight fit around the substrate by forming an extensive hydrogen bond network of interactions that stabilize dGTP for subsequent hydrolysis (Fig. 3B, C). In contrast, modeling of dTTP, dATP and dCTP binding to the active site shows that these nucleotides bind loosely in the pocket (due to their smaller size) and establish fewer hydrogen bonds with the purine or pyrimidine rings (Fig. S3H).

### GTP binding and inhibition of Ec-dGTPase activity

Given that *Ec-*dGTPase showed high specificity towards the guanosine ring, and cellular nucleotide concentrations of GTP can achieve levels 100-1000 fold higher than dGTP (36, 37), we suspected that GTP may play a role in *Ec-*dGTPase cellular regulation. To test our hypothesis, we analyzed the effects of GTP on *Ec-*dGTPase activity in the presence of 100 µM dGTP. Our results indicate that at physiologically relevant nucleotide ratios, GTP acts as a competitive inhibitor against *Ec-*dGTPase resulting in the loss of dGTP hydrolysis (Fig. 4A). To further understand this interaction, we obtained the structure of the GTP-bound enzyme to 3.25 Å by similar cross-linking and soaking methods under physiological conditions.

**Figure 4.**
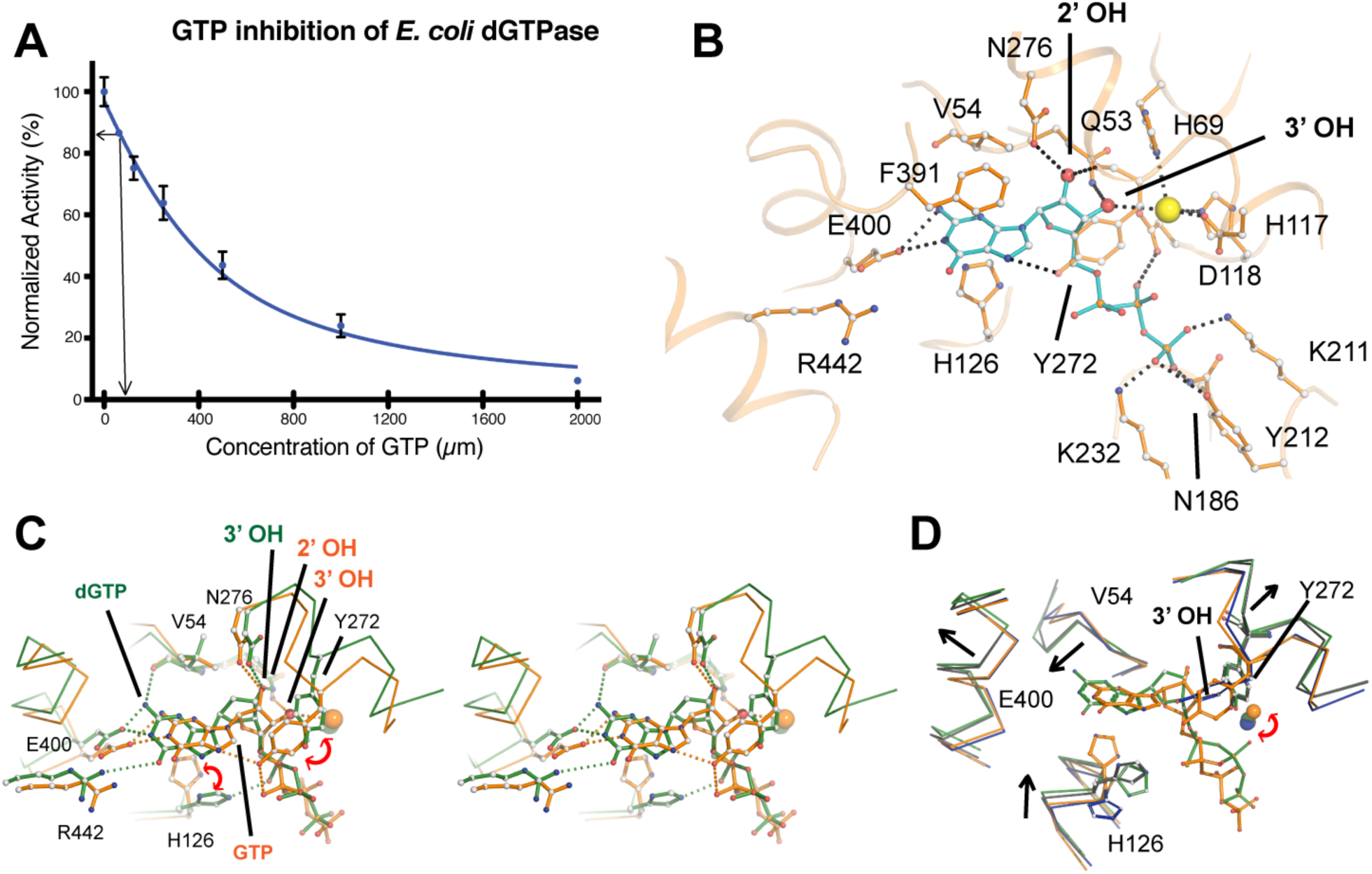
Inhibition of *Ec-*dGTPase activity by GTP. **A)** Activity of *E*c-dGTPase in the presence of increasing concentrations of GTP (µM) and 100 µM dGTP substrate. Enzymatic activity assays were repeated three times and the standard deviation plotted (n=3). The arrow indicates dGTPase activity in the presence of 100 µM GTP. **B)** Ball and stick representation and potential hydrogen bond interactions (black dashes) between GTP (cyan) and active site residues of dGTPase (orange). Asp^276^ and Gln^53^ form H-bonds with the 2′-OH and the 3′-OH, respectively. Coordination of the metal by an oxygen from the *α*- phosphate in the dGTP-bound form is swapped by the 3′-OH of the ribose, resulting in 1.5 Å displacement with respect to its position in the dGTP-bound form; as result of this positional change, HD-motif residue Asp^118^ no longer forms part of its coordination sphere. **C)** Stereo and ball and stick representations of the overlay between the dGTP-(forest green) and the GTP-bound (orange) structures illustrating that the two binding pockets differ significantly. **D)** Overlay of the substrate bound structures (dGTP and dGTP-1-thiol, green and black respectively) with the XFEL-apo and GTP-bound (inhibited) structures (blue and orange) illustrating that GTP binding “locks” the active site hindering the conformational changes observed during substrate binding (see text). The RMSD differences between active site residues for XFEL, dGTP and dGTP-1-thiol and GTP are summarized in Table S3

Analogous to the dGTP and dGTP-1-thiol structures, GTP was observed in all six active sites. Superposition of the dGTP- and GTP-bound structures revealed conformational changes to the overall architecture of the active site pocket (RMSD of 1.2 Å for 155 C*α* atoms that comprise the active site), as well as differences relative to substrate binding (Fig. 4B-C and Fig. S4A). In contrast to the dGTP bound structure, the ribose 2′-OH of GTP occupies the position of the 3′-OH (in the dGTP-bound form), and the ribose 3′-OH draws closer to the Mn^2+^ ion becoming part of its coordination sphere (Fig. 4B). As a result, GTP does not bind deeply in the pocket, losing interactions with the carbonyl of Val^54^, which allows Tyr^272^ to stack against the pyrimidine ring and reach within hydrogen bond distance of the *α*-phosphate, pulling it away from the coordination sphere of the Mn^2+^ ion (Fig-4B-C). Intriguingly, overlay of the active sites for the GTP-bound and apo- *Ec-*dGTPase structures show minimal conformational changes between the two states (RMSD of 0.39 Å, on 155 C*α* atoms comprising the active site, Fig. 4D and Fig. S4B and C). Thus, in contrast with the extensive conformational changes observed during dGTP binding (Fig. S4B), GTP does not induce a transitional “catalytic state” of the active site.

## Discussion

Despite recent *Ec-*dGTPase structures detailing the potential of ssDNA acting as an effector molecule, fundamental questions about dGTP recognition and dGTPase activity regulation have not been elucidated. The work presented herein establishes: 1) unique approaches in both XFEL and cross-linking methodologies useful to the general crystallographic community for structure determination, and 2) a comprehensive structural framework for understanding dGTPase substrate recognition and activity.

We detailed protocols for fixed-target data collection at XFEL sources in an automated fashion (Fig. 1 and Figs. S1 & S2). Notwithstanding the proven success of the injector setups for structure determination, goniometer based fixed-target approaches at the XFEL are advantageous for: 1) data collection using delicate crystals, 2) crystals in limited supply, 3) large, radiation-sensitive crystals, or 4) crystal quality screening to prepare for injector-based experiments. Our highly efficient fixed target data collection methodology was demonstrated to provide greater than 80% crystal hit rates for SFX experiments using randomly oriented micrometer-sized crystals with varying morphologies. While crystals in MCHs were mapped prior to the experiment using UV fluorescence microscopy at the home laboratory, improvements to the standard goniometer setup at the LCLS MFX instrument (38) will incorporate UV-imaging capabilities for “on-the-fly” crystal identification and mapping. This will improve the precision of crystal positioning and provide a straightforward means to fully automate fixed target SFX experiments using a variety of MCH form factors. As UV-imaging is broadly applicable to identify a wide-range of macromolecular crystals, the general crystallographic community may easily adopt this fully automated approach for multi-crystal experiments at both synchrotron and XFEL sources. This method is also compatible with new methods for *in situ* crystal growth and data collection (Martiel, Müller-Werkmeister and Cohen, in review).

The success of our approach provided the opportunity to solve a radiation, damage-free structure of the apo- *Ec-*dGTPase enzyme from a limited number of crystals (Fig. 2). The XFEL structure revealed a distinctly closer contact of Mn^2+^ coordination towards the HD motif compared to an X-ray apo-structure and two previously published structures (21). Comparisons between the apo- and the dGTP-bound structures show that conformational changes in the active site of *E*c- dGTPase allow substrate binding. Such conformational changes involve: 1) rigid body displacements that contract and expand the substrate cage to accommodate dGTP; and 2) individual residue motions to establish hydrogen bonds with the substrates. Moreover, remodeling of the binding pocket upon dGTP- but not GTP-binding reveals a “moldable” active site where conformational changes are coupled to selectivity.

The structure of *E*c-dGTPase sheds light onto the mechanism of nucleotide selectivity in dNTPases. The dNTPase activity of SAMHD1 is regulated by dGTP or GTP/dNTP binding at its primary/secondary allosteric sites, respectively, and is mediated by tetramerization (39). The active form of SAMHD1 can bind and hydrolyze all four dNTPs with similar affinities and K_cat_ (40). Structural studies have revealed the mechanism of nucleotide binding and have shown that the shape of the catalytic pocket remains nearly identical upon binding of the four dNTP substrates (12). Overlay of the *Ec-*dGTPase and SAMHD1 structures shows that in addition to sharing similar active site architectures (Fig. 5A), HD motif residues and those involved in stacking, 3′OH discrimination, contacts with the sugar moiety, and catalysis, are highly conserved (Fig. 5B). These interactions provide stabilization of the purine or pyrimidine rings and the ribose and phosphates, but do not allow nucleotide-type discrimination. Thus, whereas *Ec-*dGTPase residues at the nucleotide ring end of the binding pocket mimic Watson-Crick interactions with the amine or ketone groups of dGTP, SAMHD1 residues are separated by a 7 Å cleft from the dNTPs (Fig. 5 C-F). Indeed, previous high-resolution structures of nucleotide-bound SAMHD1 show that water molecules bridge interactions between the nucleotide ring and active site residues (PDB 4TNQ)(11), thus abolishing nucleotide type discrimination.

**Figure 5.**
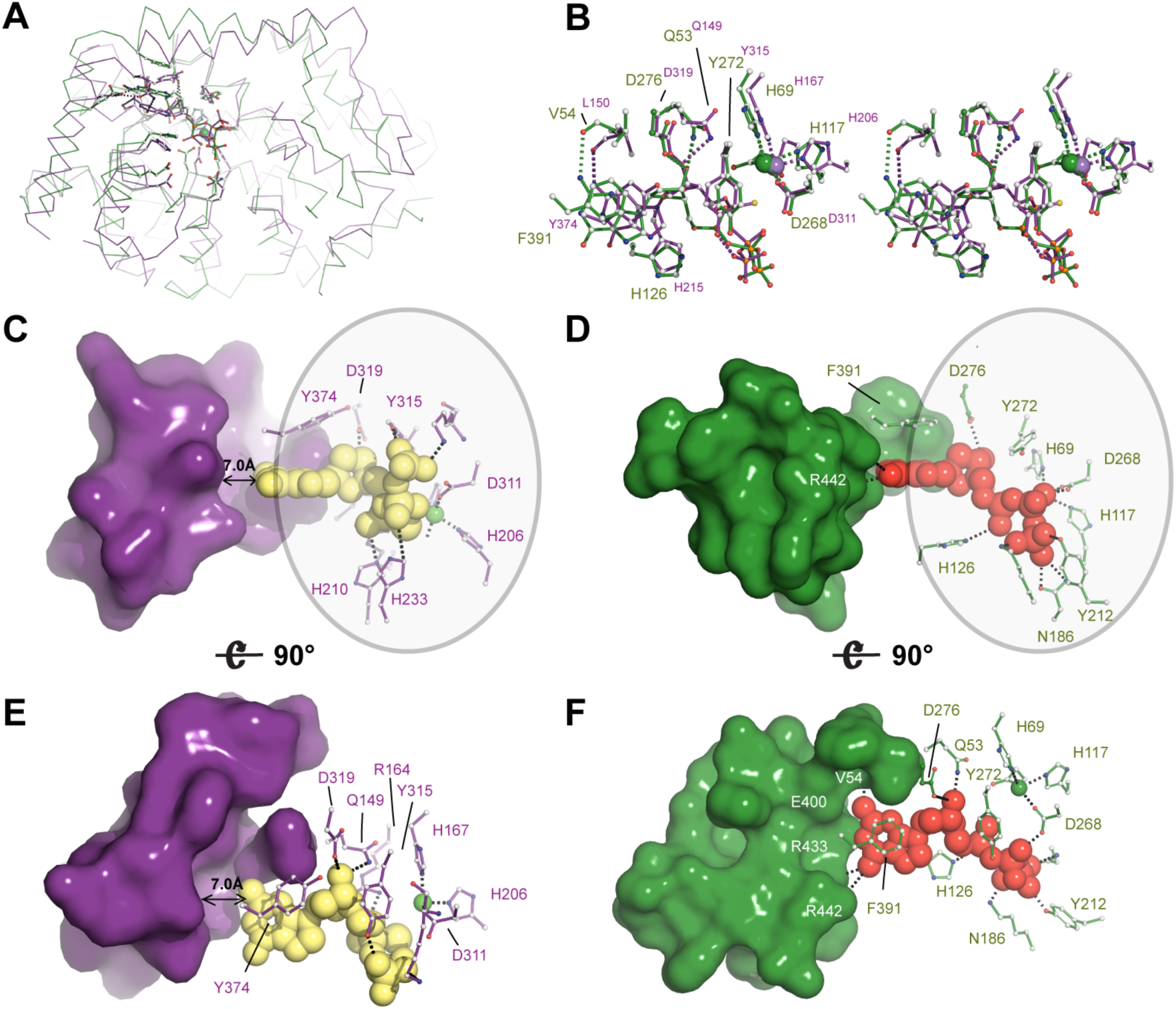
Structural and enzymatic insight into the mechanism of *Ec-*dGTPase activity. **A**) Ribbon representation of the overlay between SAMHD1 (PDB 4BZC) (purple) and *Ec-*dGTPase (forest green) illustrating fold conservation of the enzymatic cores. **B**) Stereo and ball and stick representation of the overlay between SAMHD1 and *Ec-*dGTPase active sites illustrating residue type and geometry conservation. **C) and E)** Surface and ball and stick representation of SAMHD1 active site residues illustrating that most contacts with the dNTP involve interaction with the ribose, the phosphates and the purine or pyrimidine ring (circle). No interactions with SAMHD1 residues that could confer specificity are possible since a 7 Å gap separates them from the dNTP. Thus, dNTPases bind shared motifs **D)** and **F)** A similar set of interactions takes place in dGTPase, however, dGTP selectivity occurs through formation of four hydrogen bonds.

Given that the activity of SAMHD1 is regulated by its primary/secondary allosteric sites, questions remain about the existence of an allosteric site in dGTPases. Enzymatic studies showed a 3-fold decrease in the apparent *K*_*m*_ (but had no effect on *V*_*max*_) of *Ec-*dGTPase bound to ssDNA, reflecting substrate binding enhancement (21). Moreover, the crystal structure revealed that ssDNA (found at the interface of two monomers approximately 25 Å away from the active site) triggered conformational changes affecting the catalytically essential Tyr^272^ (Fig. S2C). The structure of *Ec-*dGTPase bound to dGTP sheds light into this matter. Comparisons between the active site residues of the apo-, dGTP- and ssDNA-bound forms illustrate that the later combines features of the apo- and dGTP-bound structures (Fig. S5A). The net effect of ssDNA binding is an increase in the volume of the binding pocket by opening the tyrosine “gate” and hence improving binding efficiency, reconciling the observed three-fold decrease in *K*_*m*_.

All together, we utilized a combination of X-ray Free Electron Laser (XFEL) and chemical crosslinking methods to successfully reveal the molecular basis of *Ec-*dGTPase substrate recognition and the regulation mechanism. Together with the limited known structures of dNTPases in both prokaryotic and eukaryotic organisms, our results provide structural insight on the metabolic regulation of tightly controlled dNTP pools in DNA replication and cellular survival.

## Methods

### Cloning, Protein Purification and Crystallization

The cDNA encoding wild type *Ec*-dGTPase was a generous gift from Dr. Charles C. Richardson (Howard Medical School). It was cloned into the pET21 vector with a His-tag at the N terminus, expressed in E. coli Rosetta 2 (DE3) cultured in Luria-Bertani medium using 0.4 mM isopropylâ- D-thiogalactopyranoside for induction at 18 °C for 16 h. Proteins were first purified using a 5-ml Ni-NTA column (GE Healthcare), then followed by gel-filtration column chromatography (Hi-Load Superdex200 16/60, GE Healthcare) equilibrated with a buffer containing 25mM sodium phosphate, pH 7.5, 150 mM NaCl, 1 mM DTT, 10% glycerol, and 0.02% sodium azide.

The dGTPase crystallization solution containing 20 mM Tris-HCl, pH 8.0, 200 mM NaCl, 2 mM MnCl2, 0.02% azide, 3% Glycerol. Crystals were grown at 16°C with the sitting drop vapor diffusion method by mixture of 2 uL protein (15 mg/ml) with 2 uL crystallization buffer (100 mM Tris-HCl, pH 8.0, 1.6M AmSO_4_). Crystals were improved by dehydration in 3.5M AmSO_4_ (100 mM Tris-HCl, pH 8.0).

### Crystal growth and mounting on MCHs

Purification, assembly, and crystallization of RNA Polyermase II (Pol II), Pol II – Spt4/5 complexes was followed as previously described (38). To mount crystals onto MCHs (Fig. 1A), crystals were first cryo-protected with increasing concentrations of mother liquor. Crystal drops were increased to ~10 µL to prevent dehydration during the loading process, and MCHs were used to penetrate the drop and extract the randomly oriented crystals. For larger *Ec-*dGTPase macro-crystals (300 x 300 x 200 µm), 8 µL of cryo-solution were pipetted onto the MCH, followed by manual mounting onto the MCH via loop transfer or pipetting. To improve visualization and decrease background diffraction from solvent, excess fluid was carefully removed from MCHs using filter paper. Optimization of this step is crucial, as wicking away excess solvent may also result in the loss of crystalline sample.

### Brightfield and UV-microscopy imaging and crystal identification

Brightfield and UV microscopy was employed to identify and locate crystals in relation to the MCH reference points (Figs. S1, S2). To this end, MCHs were placed on the stage of the JANSi UVEX UV-microscope, and after focus adjustment, brightfield and UV images covering the entire area of the MCH were acquired with the nominal 5X objective using 0.1 and 1 s exposure times, respectively. After image acquisition, the crystals and MCHs were immediately flash-cooled in a liquid nitrogen bath and transferred into a SSRL cassette. A macro to detect crystals was developed using *ImageJ*, a public domain, Java-based image processing program developed by the NIH (41). *ImageJ* Macro details are described in supplemental methods.

### Post-crystallization cross-linking of E. coli dGTPase for ligand soaking

*Ec*-dGTPase crystals grown in 0.1 M Tris-HCl pH 8.0 and 1.6 M Ammonium Sulfate were transferred step-wise into 4M K/Na Phosphate pH 5.5 and 0.1 M HEPES pH 8.0. After overnight incubation, glutaraldehyde was added to the reservoir at a final concentration of 2.5%, and cross-linking proceeded for 2 hours before quenching with the addition of 0.1 M Tris-HCL pH 8.0. Cross-linked crystals were washed thoroughly with low-salt reservoir (20 mM Tris-HCl pH 8.0, 0.2 M Sodium Chloride, 2 mM Manganese Chloride, 0.02% azide, 3% glycerol) amenable to ligand-soaking, and incubated overnight with reservoir + 30% glycerol. Ligand (dGTP or dGTP- 1-thiol) with 5 mM final concentrations were added to the crystals and incubated overnight, before flash freezing in liquid nitrogen.

### Automated data collection using MCHs at the LCLS

Prior to the experiment, a reference file (Table S1) containing crystal coordinates and coordinates of four reference points for each MCH stored in a 96 sample pin storage cassette was read into the *DCS/BLU-ICE* beam line control software (42). During the day of the experiment, the SAM robot (43) was used to mount each MCH onto the beamline goniometer followed by a manual, semi-automated alignment procedure where the MCH is rotated face on to the on-axis microscope and the four reference markers are clicked in a clockwise order from within a video display of the software interface. Following this procedure, the location of the crystal coordinates is displayed over the video image of the mount (Fig. S1E) and are visually inspected. If necessary, a graphical interface enables the experimenter to remove or shift egregious crystal positions or shift the location of the reference points to improve accuracy. Updated crystal positions are stored and the user is prompted to begin automated data collection. During automated data collection, which each crystal is translated into the beam position between each X-ray pulse. This process was repeated for each MCHs in the cassette.

Diffraction experiments on *Ec-*dGTPase and Pol II complexes were done using 9.5 keV X-ray pulses with 40 fs duration and an 8 µm beam focus at the X-ray interaction point. Diffraction images were recorded on a Rayonix MX325 detector and processed using the cctbx.xfel software package (44, 45). Synchrotron based X-ray diffraction experiments of single dGTPase crystals were performed on SSRL beamline BL12-2 and the APS beamlines 22ID and 23IDD. Data were processed using XDS and SCALA software packages (46, 47). Single anomalous diffraction experiments of selenium methionine labelled dGTPase crystals were collected at 12.656 keV with inverse beam every 15° of oscillation data.

### Structure determination and refinement

Selenium substructures were determined with SHELXC/D (48), using a resolution cutoff of 4.4 Å corresponding to a CCanom = 0.301. Substructure solutions were utilized in the CRANK pipeline (49) resulting in an initial, experimentally phased structure of *Ec-*dGTPase (Fig. S5A) which was then manually built in *Coot* and refined in BUSTER. Subsequent *Ec-*dGTPase apo (XFEL), GTP-bound, dGTP-1-thiol, and dGTP-bound structures were solved by PHASER (50) using the Se-Met structure as a search model. All structures were refined using Phenix (51) and BUSTER (52), followed by several cycles of manual refinement in Coot (53). All superpositions and figures were rendered in PyMOL. Potential hydrogen bonds were assigned using a distance of <3.5 Å and an A-D-H angle of >90°, while the maximum distance allowed for a van der Waals interaction was 4.0 Å.

### Enzymatic assay of Ec-dGTPase activity

Purified wild-type and active site variants were dialyzed overnight into reaction buffer (20 mM Tris pH 7.8, 50 mM NaCl, 3% glycerol, 5 mM MgCl_2_) and concentrated to 2 mg/mL. For phosphydrolase experiments, 2µM enzyme was incubated with 100 µM dGTP (TriLink Biotech) at room temperature. Activity was monitored by quenching the reaction with 50 mM EDTA at 5, 10, 30, 60 and 120 min time-points. Quenched reactions were injected onto a C18 M column (Phenomenex) against 10 mM Ammonium Phosphate (pH 7.8) and 5% Methanol, and deoxyguanosine product was eluted with a gradient to 30% Methanol. To test the effect of GTP on enzymatic activity, enzyme was assayed in a similar manner in the presence of 100 µM dGTP and increasing concentrations of GTP (0-2 mM).

## Acknowledgements

We would like to acknowledge Michael Becker and Craig Ogata at GM/CA (Argonne National Laboratory), and Irimpan Mathews, Clyde Smith, and Ana Gonzalez at the Stanford Synchrotron Radiation Lightsource (SSRL) for their support during data collection. We thank D. Lee for home-source X-ray technical support. The cDNA of dGTPase was kindly provided by Charles C. Richardson at Harvard University. This work was supported by NIH grants R01GM116642 (J.A.) R01GM112686 (G.C.), and R01GM117126 (N.K.S.). G.C. also acknowledges support from BioXFEL-STC1231306. Use of the Linac Coherent Light Source (LCLS), Stanford Synchrotron Radiation Lightsource, SLAC National Accelerator Laboratory, is supported by the U.S. Department of Energy, Office of Science, Office of Basic Energy Sciences under contract no. DE-AC02-76SF00515. The SSRL Structural Molecular Biology Program is supported by the DOE Office of Biological and Environmental Research and by the National Institutes of Health, National Institute of General Medical Sciences (including P41GM103393). The contents of this publication are solely the responsibility of the authors and do not necessarily represent the official views of NIGMS or NIH.

## Author Contributions

C.O.B and Y.W. contributed to this work equally. C.O.B., Y.W., J.A., A.E.C., and G.C., wrote the manuscript. All authors commented and approved the manuscript.

## Supplemental Methods

### Design and Fabrication of Multi-Crystal Holders (MCHs)

MCHs were developed to meet the following requirements: 1) minimal UV-background, 2) a larger size than commercially available mounts to hold substantial amounts of crystals, and 3) compatibility with the SAM robot system for robotic exchange during the diffraction experiment. Initial MCH prototypes were fabricated at the University of Pittsburgh, Swanson School of Engineering using a commercially available Stereo Lithography Machine, a 3D Systems Viper High Resolution SLA and Somos 11122XC Resin. The pointed end of the diamond shaped MCH was necessary to penetrate a crystal containing drop and mount crystals without drop displacement. Despite the success of the first-generation 3D-printed MCHs, subsequent generations of MCHs were manufactured by micron laser technology (Oregon, USA) from Mylar sheets (McMaster Carr, IL, USA) of 50 µm thickness (Fig. S1A, B). The laser-cut MCHs had better defined reference features (fiducial marks for crystal positioning), and were resilient to the physical manipulation during the crystal loading process. MCHs were affixed using epoxy to the end of a standard Hampton Research base-pin assembly for robotic exchange (Fig. S1B).

### Macro parameters for detecting crystals on MCHs

UV microscopy images were loaded into *ImageJ* and brightness/contrast features were adjusted to highlight UV positive regions (Fig. 1B and Figs. S1, S2). Images were subsequently processed using a macro developed in our laboratory. Sample input parameters for our macro are listed below:

**run**(“Smooth”);

**run**(“Threshold”, “method=Huang ignore_black white setthreshold”);

**setThreshold**(40, 900);

**setOption**(“BlackBackground”, false);

**run**(“Erode”);

**run**(“Analyze Particles…”, “size=150-10000 circularity=0.25-1.00 show=Outlines display summarize in situ”);

Modifications to brightness threshold, particle size, and circularity are critical to the selection of bright areas that indicate UV-visualized crystals, which are converted into crystal profiles (Fig. S2C) and a list specifying crystal size and centroid position (in x-y jpeg pixel coordinates, see Table S1). The pixel coordinates of the four MCH reference points were identified in relation to the crystal coordinates (Fig. 1B and Fig. S1E, red dots) through examination of a corresponding brightfield image (Figs. S1C and S2A, D). For larger crystals (Fig. S2D-F), a library of masks was created to define spacing along a crystal to translate into unexposed crystal volumes. To utilize this feature, the macro was altered to allow for the overlay of the mask as follows:

**run**(“Subtract…”, “value=50”);

**run**(“Smooth”);

**run**(“Threshold”, “method=Huang ignore_black white setthreshold”);

**setThreshold**(30, 255);

**setOption**(“BlackBackground”, false);

**run**(“Make Binary”, “thresholded remaining”);

The resulting image was then overlaid with the mask (Fig. S2E, inset) followed by:

**run**(“Make Binary”, “thresholded remaining”);

**run**(“Analyze Particles…”, “size=40-2000 circularity=0-1.00 show=Outlines display summarize record in situ”);

The minimum size was set to the same size (in pixels) as the mask size and circularity began at 0 since the mask creates square shapes. Coordinates for each distinct beam position were saved with an angular offset for helical data collection.

## Supplementary Figures and Legends

**Supplementary Figure 1.**
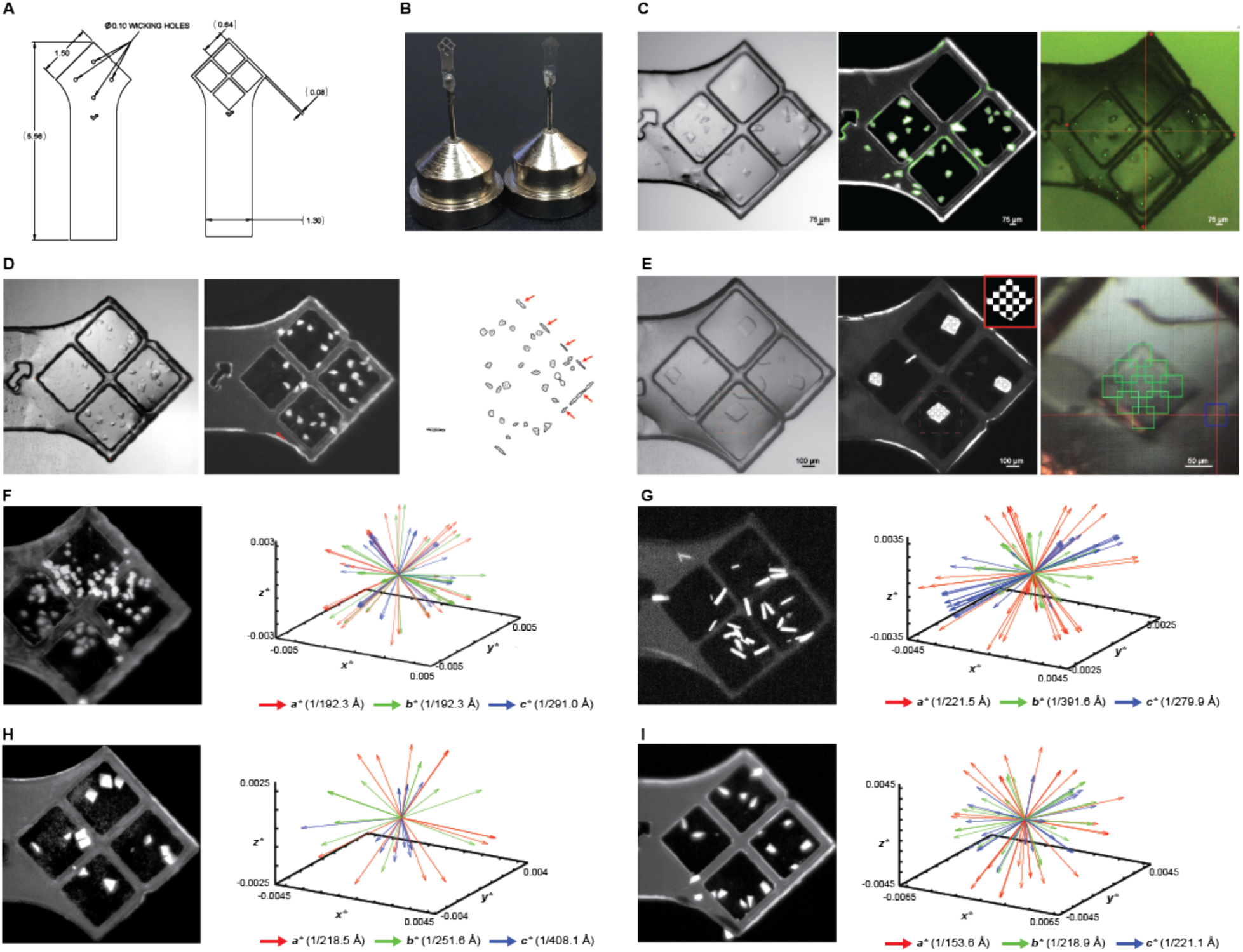
Design and identification of crystals mounted on Multi Crystal Holders. **A)** AutoCAD DXF schematic illustrating the dimensions of MCHs for laser printing onto Mylar sheets. **B)** Assembly of MCHs achieved by epoxying laser-printed MCHs onto the pin-base aperture of a Hampton loop. **C)** Left and Middle Panels: Brightfield and UV fluorescence microscopy images of Pol-Spt4/5-DNA mounted crystals. Green outlines indicate the crystal positions determined after particle analysis in ImageJ. Right panel: *BluIce* Image of Pol-Spt4/5 crystals mounted at XPP endstation. Coordinates generated in ImageJ (see Table S1) were uploaded, and after identification of reference fiducial marks (red asterisks) and beam positions were assigned to each crystal (green boxes). **D)** Brightfield image, UV fluorescence image, and crystal positions as revealed by ImageJ. Coordinates for incorrectly identified UV fluorescence signal on the edge of the MCH (red arrows, right panel) are manually removed from the final coordinate list before data collection. **E)** Large crystals of the Pol II – TFIIB – DNA complex illustrate the multi-shot strategy with user specified spacing along the crystals surface. **F-I)** Reciprocal space representation of the basis vectors of indexed crystals that were singularly exposed for **F)** *Ec-*dGTPase, **G)** RNA Polymerase II T834P variant, **H)** RNA Polymerase II – TFIIB – DNA, and **I)** RNA Polymerase II – Spt4/5 – DNA crystals.

**Supplementary Figure 2.**
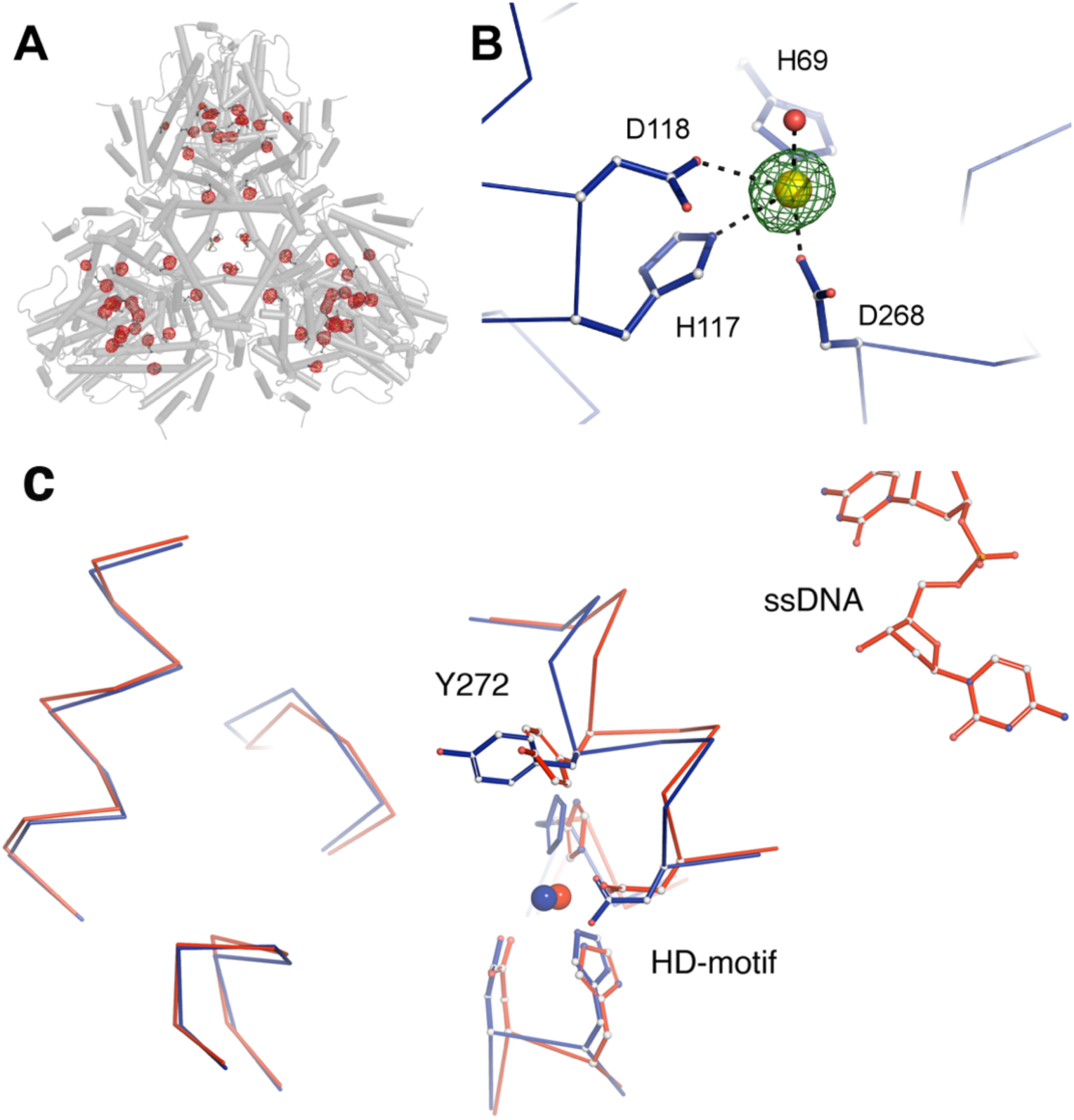
Se-Met substructure and comparison of active site residues in apo- *Ec-*dGTPase structures. **A)** Tube representation illustrating the hexameric *Ec-*dGTPase structure determined by experimental phasing from selenium methionine labeled protein. The Se anomalous difference map is shown in red, contoured at 8*σ*. **B)** The Fo-Fc map countered at 6*σ* of the apo-XFEL structure calculated after removal of Mn^2+^ ions from the refined structure (modeled Mn^2+^ ion shown in yellow). **C)** Overlay of the *Ec-*dGTPase apo-XFEL (blue) and ssDNA (red, PDB 4×9E) structures illustrating conformational differences mainly in the *α*10 helix involving Tyr^272^ due to the presence of the ssDNA located 25 Å away from the binding pocket.

**Supplementary Figure 3.**
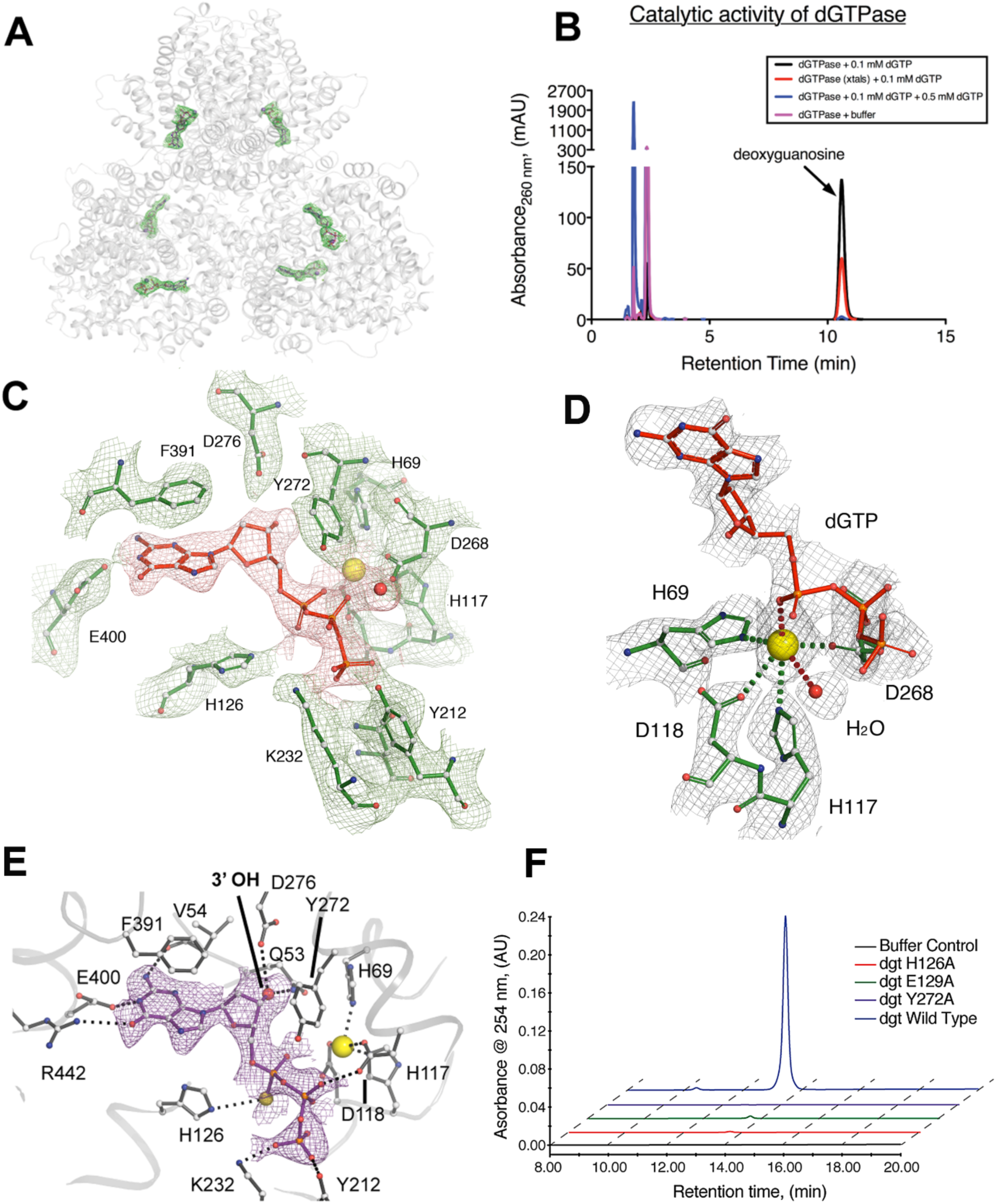
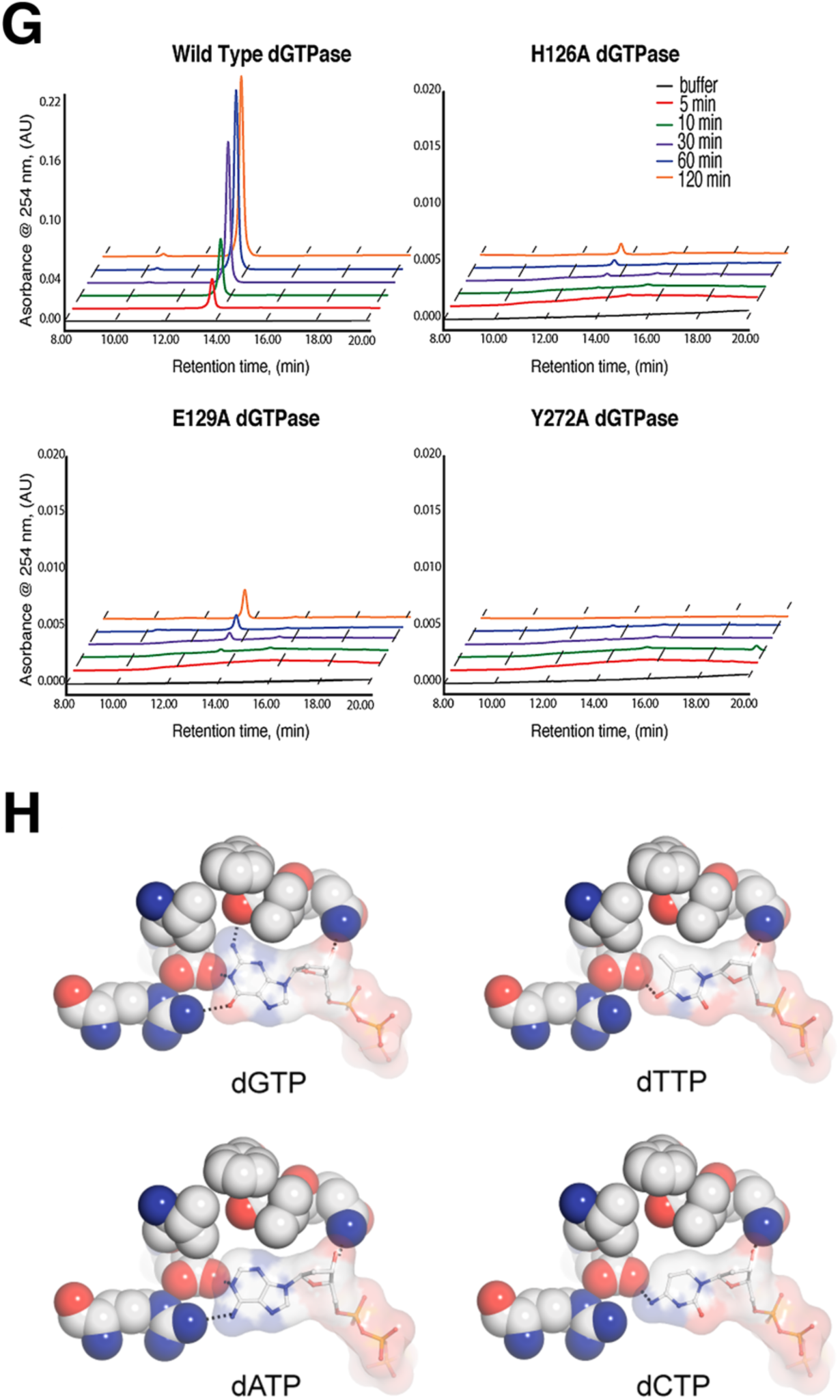
Nucleotide substrate binding, specificity, and *Ec-*dTPase activity. **A)** Fo-Fc map countered at 3*σ* showing electron density corresponding to dGTP. The presence of the substrate was observed in every pocket of the hexamer (modeled dGTP shown). **B)** Catalytic activity of *Ec-*dGTPase crystals in the presence of 100 µM dGTP, illustrating that cross-linked crystals (red line) are still catalytically active when compared to wild-type (black line) or *Ec-* dGTPase crystals in the presence of 0.5 mM thio-GTP (blue line). **C)** Final refined 2Fo-Fc map countered at 1.5 *σ* after applying negative B-factor sharpening =150. The quality of the map allowed full tracing of the structure. **D)** Final refined *2F*_*obs*_ *– F*_*calc*_ map contoured at 2.0 *σ* and ball and stick representation of Mn^2+^ icosahedral coordination by HD residues (forest green), dGTP and a water molecule (W1) are shown as ball and stick model and red sphere respectively. **E.)**Ball and stick representation of residues involved in thio-dGTP binding and final refined *2F*_*obs*_ *– F*_*calc*_ map around thio-DGTP contoured at 1.2*σ*. The overall and active site R.M.S.D. between the two substrates bound structures thio-dGTP and dGTP is 0.4 Å and 0.3 Å respectively. **F)**Enzymatic activity analysis after 2-hour incubation at room temperature in the presence of 100 µM dGTP substrate was performed by monitoring total deoxyguanosine product using reverse-phase chromatography for wild-type (blue), Y272A (purple), E129A (green) and H126A (red) dGTPase enzymes. The representative chromatogram indicates that the activity of dGTPase mutants is significantly decreased when compared to wild-type enzyme. **G)**Enzymatic activity of individual dGTPase constructs. Representative reverse-phase HPLC chromatograms illustrating the amount of deoxyguanosine (dG) product attained after 5 (red), 10 (green), 30 (purple), 60 (blue), and 120 (orange) minutes of incubation at room temperature in the presence of 100 µm dGTP for wild-type, H126A mutant, E129A mutant, and Y272A mutant constructs. Wild-type enzyme is capable of producing an order of magnitude more dG product after 5 minutes (red line) than the mutant constructs after 120 minutes incubation (orange lines). **H)**Comparison of dGTP with models of dTTP, dATP and dCTP binding to illustrate that active site residues can form four hydrogen bonds with the amide and ketone groups of dGTP but not with the other NTPs where one interaction is observed at best.

**Supplementary Figure 4.**
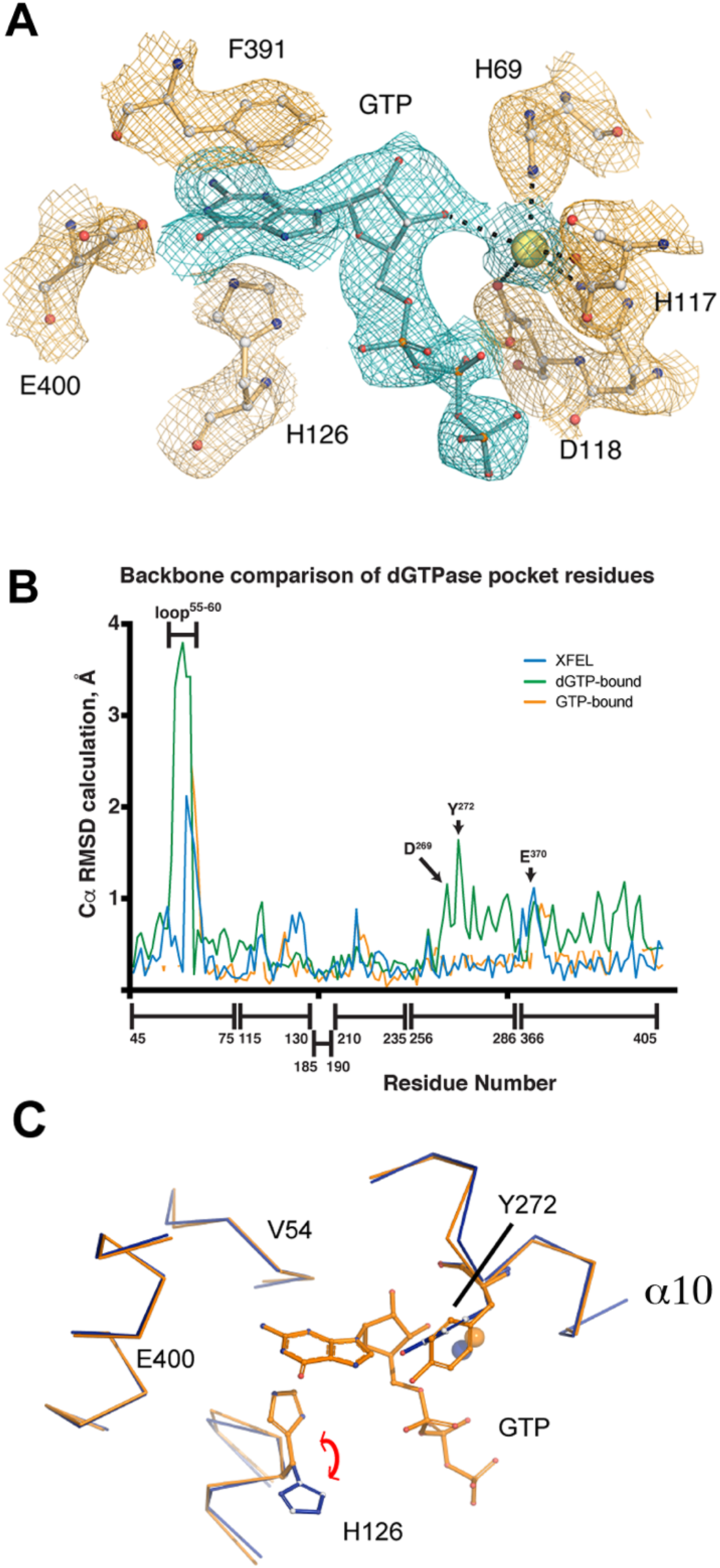
Structural differences related to GTP inhibition of *Ec-*dGTPase. **A**) Final refined *2F*_*obs*_ *– F*_*calc*_ of GTP bound to the active site pocket contoured at 1.2*σ* after applying negative B-factor sharpening=160. **B.** Graphical representation of C*α* RMSD illustrating: 1) the similarities between the apo and GTP structures (orange and blue traces) and 2) the conformational changes of the binding pocket residues triggered by dGTP binding (green trace). Positional differences for C*α*s of Val^54^, Asp^268^, Tyr^272^ and Glu^400^. The position of the catalytic His^126^ varies among the three structures illustrating its conformational flexibility. **C)** Overlay between the apo and GTP-bound structures showing the similar conformations between the two of them.

**Supplementary Figure 5.**
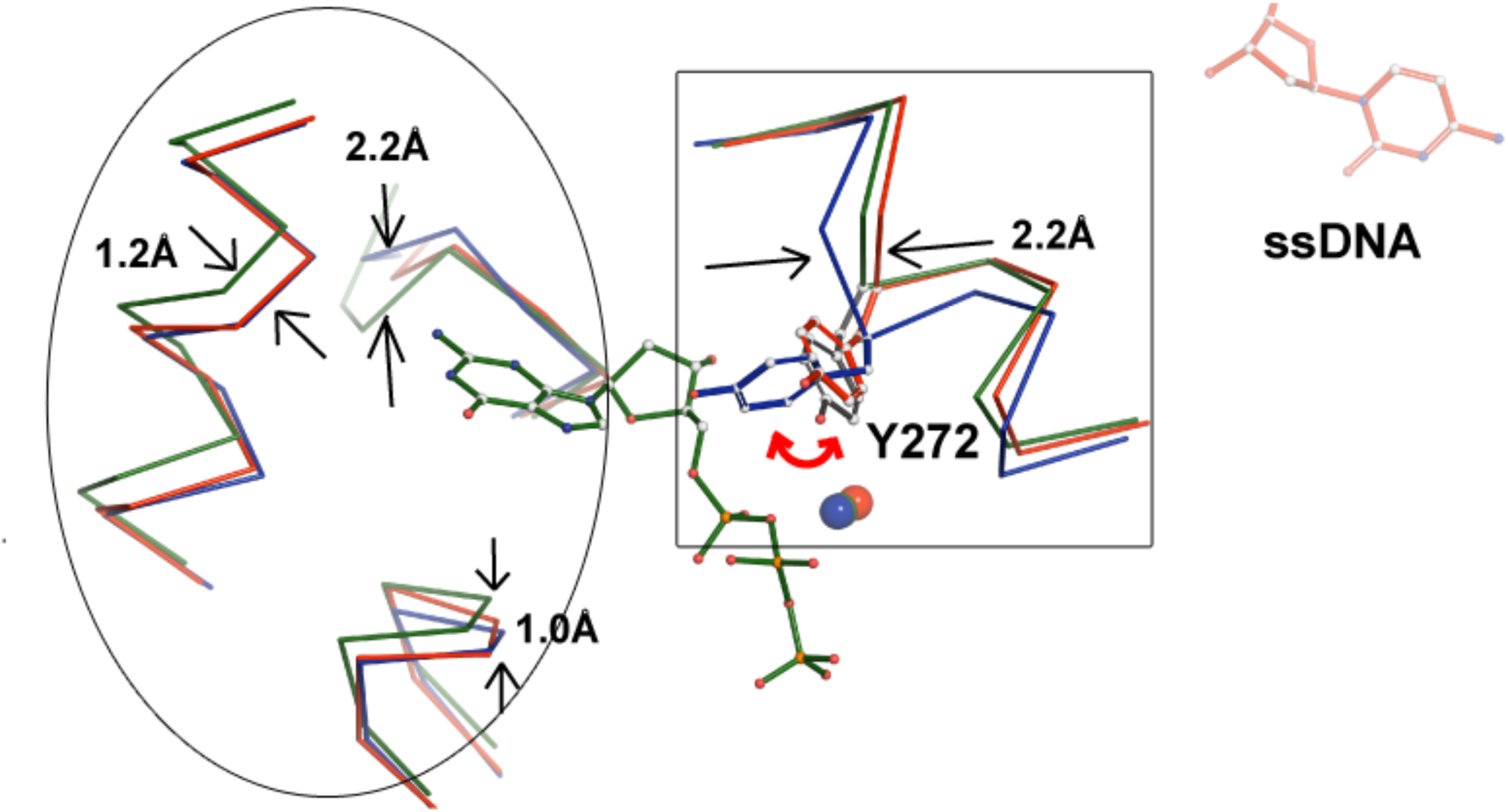
Overlay between *Ec-*dGTPase apo-structure bound to ssDNA (PDB 4×9E, red wire and red ball and stick model) with the apo-XFEL (blue wire) and the dGTP- bound structures (forest green wire). The ssDNA structure combines apo-like (overlapped red and blue traces, left ellipse) and substrate-bound like (overlapped green and red traces, right square) features. These conformational changes lead to a 20% increase in the size of the substrate binding pocket. The RMSD for C*α*s between the structures is indicated between arrows.

## Supplemental Tables

**Table S1.** Example of crystal coordinates and fiducial marks derived from ImageJ for MCH represented in Figure 1.

| *Fiducial Coordinates* | *X-Coordinate* | *Y-Coordinate* | *Angular Offset<sup>a</sup>* |
| --- | --- | --- | --- |
| Position 1 | 844 | 36 | 0 |
| Position 2 | 1430 | 690 | 0 |
| Position 3 | 794 | 1258 | 0 |
| Position 4 | 240 | 644 | 0 |

| *Crystal Coordinates* | *X-Coordinate* | *Y-Coordinate* | *Angular Offset* |
| --- | --- | --- | --- |
| Crystal 1 | 672.8 | 217.9 | 0 |
| Crystal 2 | 1045.2 | 258.0 | 0 |
| Crystal 3 | 910.4 | 259.7 | 0 |
| Crystal 4 | 397.9 | 310.3 | 0 |
| Crystal 5 | 965.8 | 322.4 | 0 |
| Crystal 6 | 616.2 | 342.1 | 0 |
| Crystal 7 | 638.1 | 379.0 | 0 |
| Crystal 8 | 1078.2 | 383.8 | 0 |
| Crystal 9 | 747.7 | 413.5 | 0 |
| Crystal 10 | 939.3 | 426.6 | 0 |
| Crystal 11 | 1207.8 | 438.5 | 0 |
| Crystal 12 | 298.2 | 438.0 | 0 |
| Crystal 13 | 534.2 | 453.5 | 0 |
| Crystal 14 | 328.4 | 480.2 | 0 |
| Crystal 15 | 804.5 | 516.6 | 0 |
| Crystal 16 | 761.4 | 531.3 | 0 |
| Crystal 17 | 215.7 | 529.6 | 0 |
| Crystal 18 | 652.6 | 560.7 | 0 |
| Crystal 19 | 384.0 | 559.1 | 0 |
| Crystal 20 | 977.7 | 607.3 | 0 |
| Crystal 21 | 506.4 | 624.8 | 0 |
| Crystal 22 | 360.8 | 621.6 | 0 |
| Crystal 23 | 817.9 | 642.7 | 0 |
| Crystal 24 | 324.7 | 710.5 | 0 |
| Crystal 25 | 452.9 | 731.0 | 0 |
| Crystal 26 | 1046.7 | 759.6 | 0 |
| Crystal 27 | 841.8 | 746.8 | 0 |
| Crystal 28 | 250.9 | 784.2 | 0 |
| Crystal 29 | 696.7 | 816.8 | 0 |
| Crystal 30 | 1226.5 | 812.9 | 0 |
| Crystal 31 | 922.8 | 839.8 | 0 |
| Crystal 32 | 411.3 | 862.5 | 0 |
| Crystal 33 | 589.0 | 893.2 | 0 |
| Crystal 34 | 1113.9 | 911.4 | 0 |

<sup>a</sup>For crystals capable of multiple exposures, and angular offset can be input to improve data completeness.

**Table S2.**
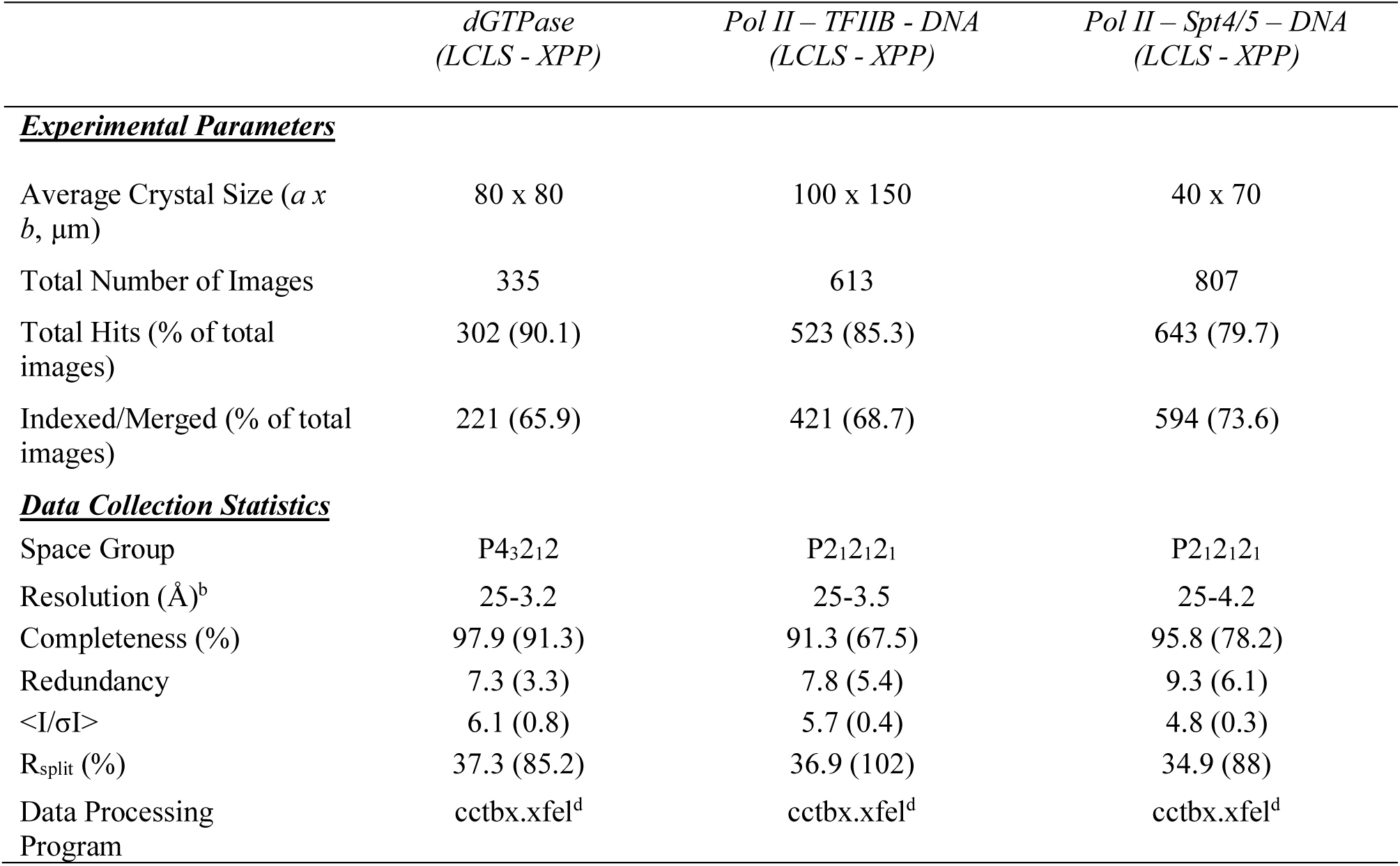
Data collection statistics and efficiency using MCHs

**Table S3.**
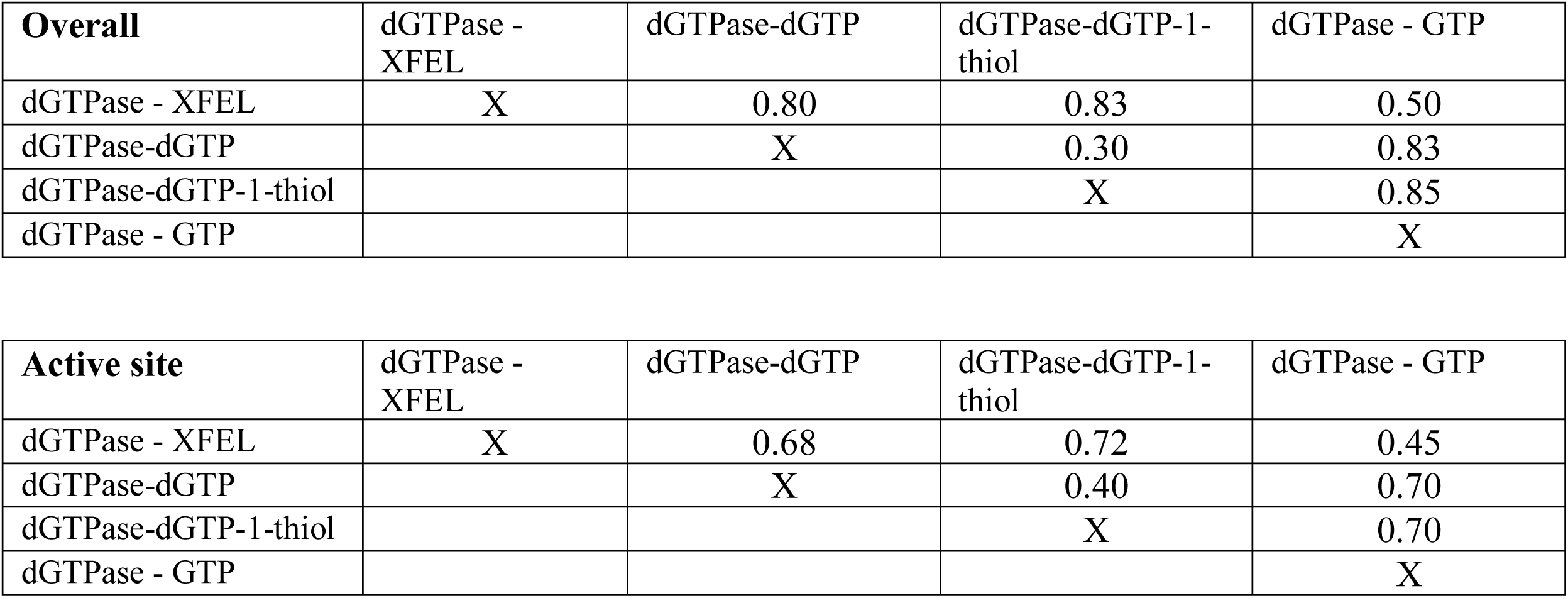
Structural overall and active site comparison R.M.S.D (Å) statistics

